# *TidyMass*: An Object-oriented Reproducible Analysis Framework for LC-MS Data

**DOI:** 10.1101/2022.03.15.484499

**Authors:** Xiaotao Shen, Hong Yan, Chuchu Wang, Peng Gao, Caroline H. Johnson, Michael P. Snyder

**Author notes:** Corresponding authors: Caroline H. Johnson and Michael P. Snyder. These authors contributed equally: Xiaotao Shen, Hong Yan, and Chuchu Wang.

## Abstract

Reproducibility and transparency have been longstanding but significant problems for the metabolomics field. Here, we present the tidyMass project (https://www.tidymass.org/), a comprehensive computational framework that can achieve the shareable and reproducible workflow needs of data processing and analysis for LC-MS-based untargeted metabolomics. TidyMass was designed based on the following strategies to address the limitations of current tools: 1) Cross-platform utility. TidyMass can be installed on all platforms; 2) Uniformity, shareability, traceability, and reproducibility. A uniform data format has been developed, specifically designed to store and manage processed metabolomics data and processing parameters, making it possible to trace the prior analysis steps and parameters; 3) Flexibility and extensibility. The modular architecture makes tidyMass a highly flexible and extensible tool, so other users can improve it and integrate it with their own pipeline easily.

---

To date, liquid chromatography-mass spectrometry (LC-MS)-based untargeted metabolomics has been proven to be an important tool in environmental, nutrition, and biomedicine research^1^. A typical full workflow for LC-MS-based untargeted metabolomics includes sample collection, data acquisition, data analysis, and biological interpretation^2^ (**Fig. S1**). Processing and analyzing high-dimensional metabolomics datasets are challenging, requiring the optimization of multiple steps such as raw data processing, data cleaning, data quality control and assessment, metabolite annotation, statistical analysis, and biological function mining^3^.

To overcome the challenges of processing and analyzing metabolomics data, the community has developed numerous tools^4,5^. However, limitations still exist. Commercial tools are expensive and only work on the associated instrument platform, online/GUI tools are user-friendly but cannot take the advantage of the cluster and server computational resources making them impractical for large-scale datasets, open-source tools typically follow limited parts of the whole bioinformatics workflow and have no uniform, specific and traceable format for data input, resulting in a complicated and time-consuming process to prepare data. In addition, different tools with different design concepts and based on different computational platforms make data sharing and reproducible analyses extremely challenging.

Here, we proposed the tidyMass project, an ecosystem of R packages that share an underlying design philosophy, grammar, and data format, which provides a comprehensive, reproducible, and object-oriented computational framework.

We first designed a specific uniform data format (“mass_dataset”) to efficiently store and manage processed untargeted metabolomics data (**Fig. 1**). In the “mass_dataset” class, the expression dataset, metadata of samples and variables are included. Additionally, the datasets in it are automatically synchronous, so when the users operate one component, it will automatically propagate the operations across all corresponding components (**Fig. S2**). This makes it easy to manipulate and maintain the consistency of the data. All the functions in tidyMass use the “mass_dataset” as their primary input data format, therefore one data format can be used for all processing and analysis steps (**Fig. 2**). Additionally, the “mass_dataset” class supports popular tools from other packages, in particular tidyverse, which is one of the most widely used tools for data science in the R environment^6^ (**Fig. S3**). This design makes the code of tidyMass more universal and straightforward, which benefits new users as they do not need to adopt new functions. Furthermore, all the parameters for the processing and analysis are stored in the “mass_data” class object, which makes it feasible to trace the prior steps and parameters (**Fig. S4**). Briefly, the “mass_dataset” class provides a simple way to manage and process metabolomics data which sets the foundation for the highly reproducible, robust, and extendable analytical framework.

**Fig. 1.**
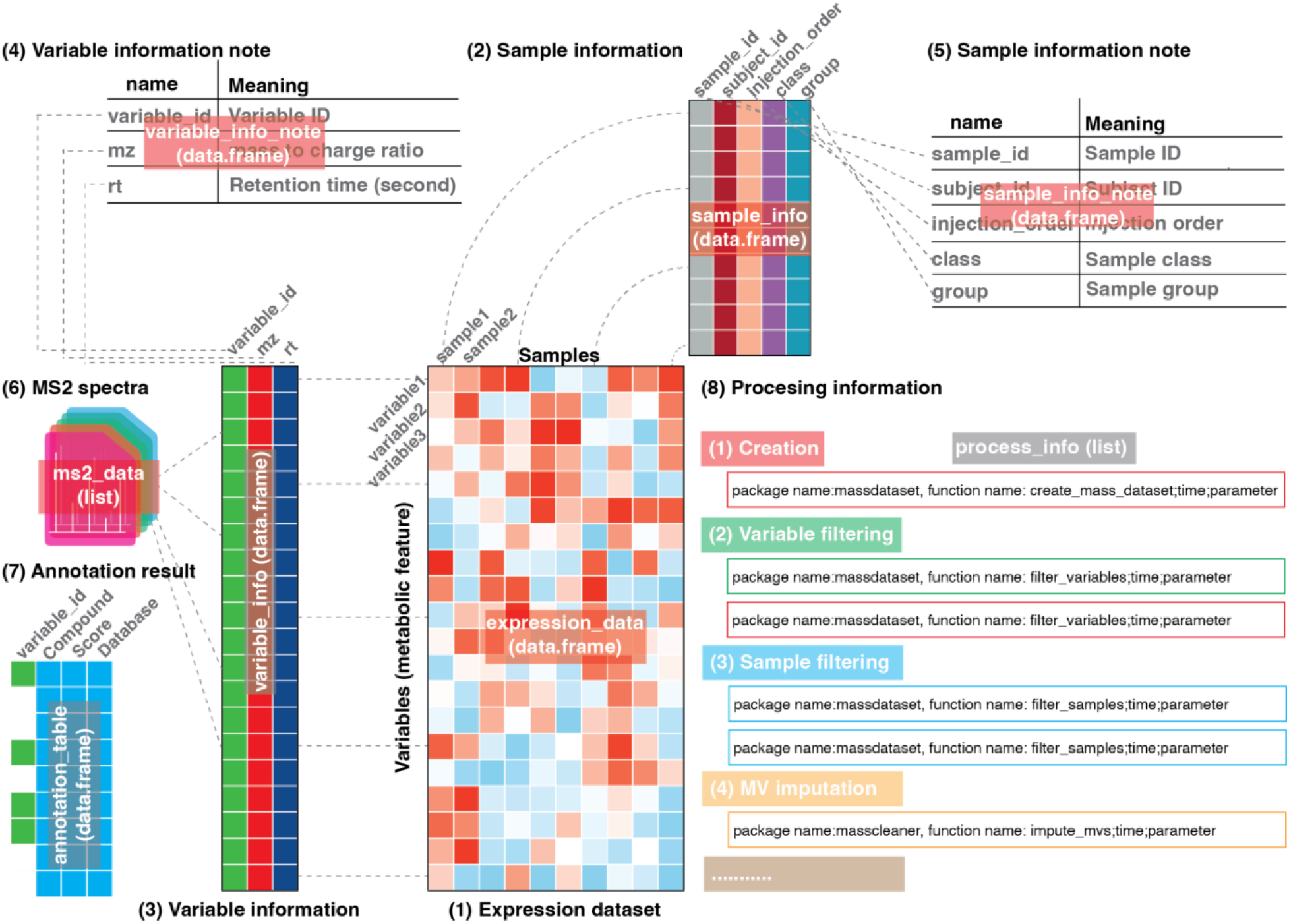
The “mass_dataset” class and its property. The “mass_dataset” class is a uniform data format, which is specifically designed for representing metabolomics data. Most functions in the tidyMass expect this class as their input format, and all the parameters for the functions can be stored in it.

**Fig. 2.**
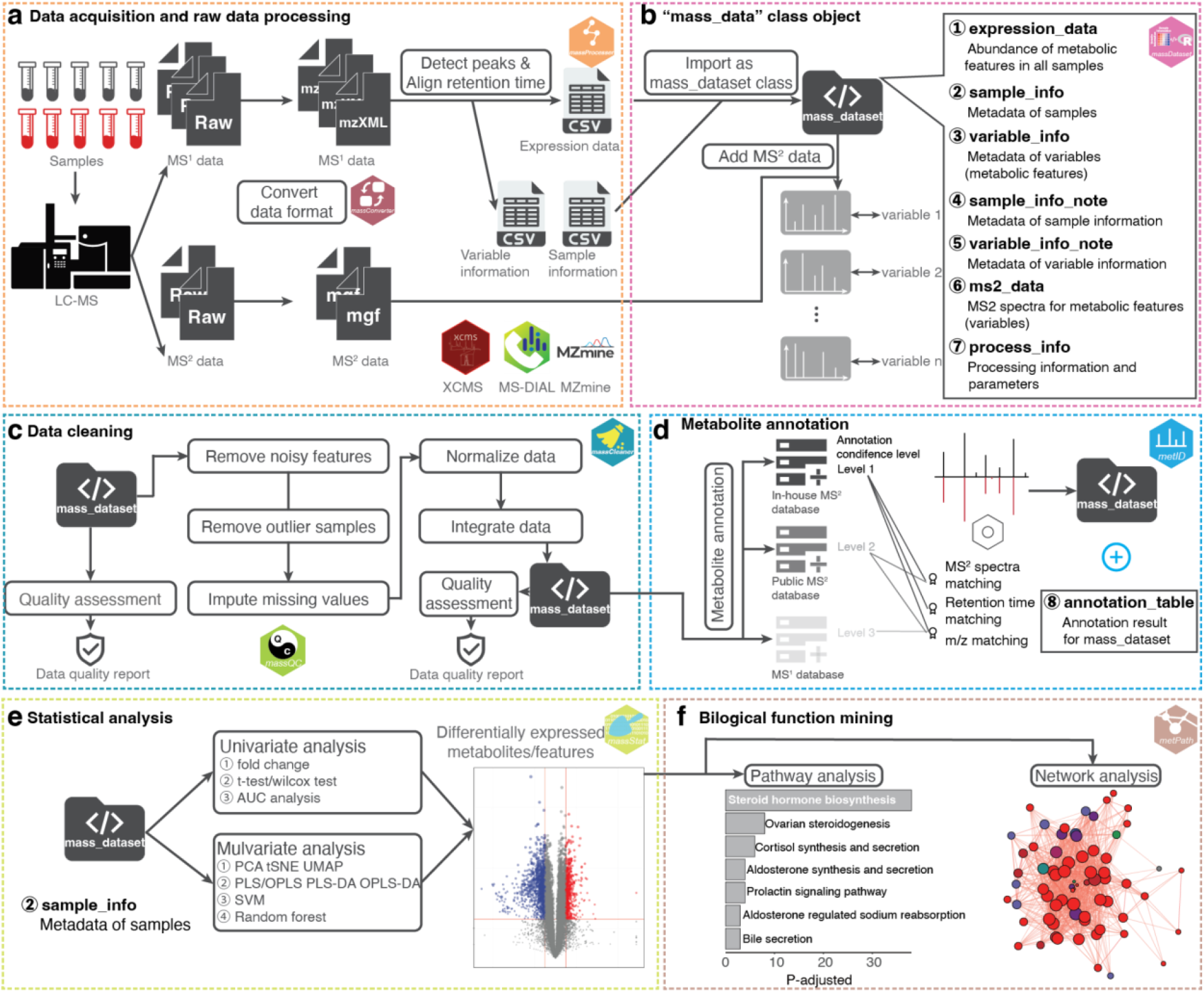
Analysis workflow of tidyMass. **(a)** Raw data processing using massConverter and massProcesser. **(b)** The “mass_dataset” class provides a uniform data format and object-oriented workflow (massDataset). **(c)** Data cleaning using massCleaner. **(d)** Metabolite annotation using metID. **(e)** Statistical analysis using massStat. **(f)** Biological function mining using metPath.

TidyMass provides a set of functions that takes the “mass_dataset” class as the input data format to perform the whole workflow (**Fig. 2** and **Table S1**). Similar to the concept of tidyverse^6^, tidyMass does not include all the functions in one package, which is flexible to both users and project managers. TidyMass is a collection of multiple R packages, where the different packages correspond to different steps of the workflow (**Fig. 2**). The modular design makes it easy for the user to find appropriate functions, and for developers to debug and extend it^7^. Briefly, the workflow begins from the package massConverter, which converts MS raw data from different vendors to other formats (**Fig. 2a**). MassConverter depends on the docker version of msconvert^8^, making it possible to use it on all computational platforms. So the data conversation can also be integrated with other processing and analysis steps in one code script, which makes the end-to-end reproducible analysis possible. Next, raw data processing, peak picking and grouping are performed by the massProcesser package based on XCMS^9^, an object (“mass_dataset” class) is generated for subsequent analysis in this step. Before moving forward to statistical analysis, data cleaning is performed to remove unwanted variation by the massCleaner package^10^, which carries out noisy features and outlier sample removal, missing value imputation, data normalization and integration. In the next step, the metID package performs metabolite annotation using in-house or public databases^11^. All the statistical analyses are aimed at finding the potential differentially expressed metabolites using the massStat package^12^. Finally, pathway enrichment analysis is implemented to identify biological functions using the metPath package. Notably, in any step of the workflow, the massQC package can be used to assess the data quality.

Data sharing and reproducible analysis are of utmost importance to avoid biased findings^13^. Unfortunately, reproducibility and transparency for metabolomics within the R environment are less satisfactory than for other types of omics data. Multiple tools offer different parameters, options, and output formats for users. TidyMass is designed to achieve reproducibility and transparency by two aspects. First, the object-oriented class makes it easy to share the data and trace the processing information^14^. Second, with the uniform data format and modular design, the users can seamlessly combine all the processing and analyzing steps in an integrative manner in one code script (*e.g.*, Rmarkdown, notebook). In addition, all the steps are optional and the order of execution is customizable, which means that the users can create and optimize customized sharable and reproducible pipelines based on their experimental design and aims. Furthermore, as docker technology is more and more popular in reproducible analysis, we also provide a docker version of tidyMass, containing a R/Rstudio environment and all the tidyMass packages, which makes it possible for users to share all code, data, and even analysis environment based on tidyMass.

To demonstrate the application of tidyMass for the processing of metabolomics data, we used data from colorectal cancer (CRC) patient tissues to identify metabolites of CRC by sex of the patient^15^ (**Table S3, Fig. S5**). First, raw data were converted to mzML format through ProteoWizard^8^, followed by massProcessor to extract the metabolic features. Features with more than 20% missing values (MV) in QC samples or more than 50% MVs in all the study groups were considered as noisy features and were removed. K-nearest neighbors (KNN) was applied for MV imputation, and support vector regression (SVR) enabled data normalization using massCleaner (**Fig. S6**). For metabolite annotation, two in-house databases were constructed using metID, that contain 71 and 55 metabolites in HILIC and RPLC modes, respectively. The databases contain the accurate mass and experimental retentional time of metabolites. A public database^11^ was also used for metabolite annotation. Finally, the redundant annotations were removed based on the annotation score^11^, and 74 metabolites were identified using the in-house database and up to metabolomics standards initiative (MSI) level 2^16^. Only the annotations with level 2 were used for subsequent analysis. We then detected the differentially expressed metabolites between tumor tissues compared to normal controls for males and females separately, using massStat (**Fig. S7, Fig. 3a**). Furthermore, metPath was used for pathway enrichment. In addition to our previous findings wherein sex-related differences were observed in methionine, polyamine, pentose phosphate pathways, methionine metabolism and polyamine metabolism^15^, we also observed differential enrichment of additional pathways in tumors from female and male patients (**Fig. 3**). For example, ferroptosis and bile acid synthesis was only enriched in tumors from male patients (**Fig. 3 b**). Glutathione metabolism, the cAMP signaling pathway, cGMP-PKG signaling pathway were all enriched in tumors from female patients, but not from males. In addition, tidyMass expedited the analytical workflow, making it more straightforward to analyze and reproducible using a code script (**Supplementary Data 1** and **2**). Additionally, a docker image containing the data, code and analysis environment is also provided for more straightforward reproducible analysis (https://hub.docker.com/r/jaspershen/tidymass-case-study).

**Fig. 3.**
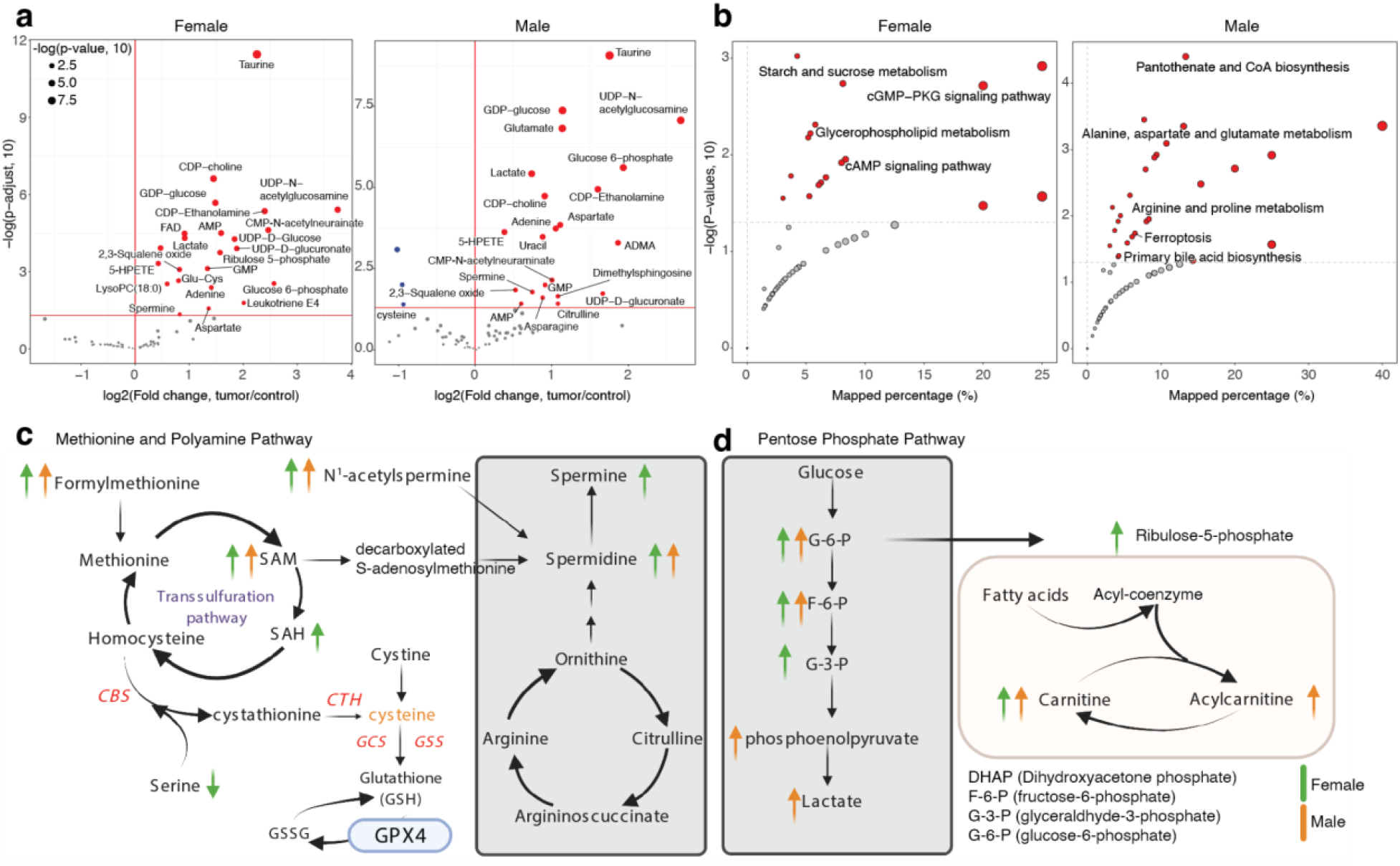
Biological function mining for the case study. **(a)** Volcano plots to show the differentially expressed metabolites. **(b)** Pathway enrichment analysis. Only the sex difference pathways were labeled. Sex differences in **(c)** methionine and polyamine pathways, and **(d)** pentose phosphate pathway metabolism.

In summary, the tidyMass project is an ecosystem of R packages that share an underlying design philosophy, grammar, and uniform data format, which provides a comprehensive, transparent, reproducible, and object-oriented computational framework for LC-MS-based metabolomics data processing and analysis within the R environment. As such, a complete website for tidyMass is publicly available (https://www.tidymass.org/). TidyMass can provide great benefit for the metabolomics field, particularly in the two following aspects. 1) Data sharing, tracing, and reproducible analyses. TidyMass provides a specific uniform data format and a whole object-oriented workflow, including a docker image, making data sharing, tracing, and reproducible analysis more straightforward, providing metabolomics researchers the ability to share and repeat analysis feasibly. 2) Flexibility and extensibility. The object-oriented and modular design concept allows for the easy integration of other tools with tidyMass, therefore making tidyMass flexible and extensible within the metabolomics community. An example that illustrates the functions of the tidyverse can be located here: https://massdataset.tidymass.org/articles/tidyverse_verse. However, as a fast-growing field, some widely used metabolomics tools are not wrapped up or supported in tidyMass, such as GNPS^17^ and MetaboAnalystR^18^, it is capable to convert the “mass_dataset” class to the eligible data format for these tools in the future. Meanwhile, as an open-source tool, tidyMass can be easily implemented into the other pipelines.

## Methods

### TidyMass project

TidyMass project is an ecosystem of R packages that share an underlying design philosophy, grammar, and data structure, which utilizes the concept of tidyverse^19^. To address the challenges of data sharing, reproducible analysis, and extensibility, we adopted the object-oriented and modular design concepts, which are also leveraged by other tools^7,14^.

#### Object-oriented workflow

In tidyMass, the “mass_dataset” class is designed specifically for storing the metabolomics data and relevant metadata. Most of the functions in all the packages use it as the input data and output format. Based on the “mass_dataset” class and the pipeline function (%>%) from tidyverse, tidyMass provides an object-oriented workflow of data processing and analysis, which is clear and more straightforward.

#### Modular design

In tidyMass, different packages correspond to different steps of the whole workflow for LC-MS-based untargeted metabolomics data processing and analysis. The fundamental functions are placed in one package named massTools, therefore all the other packages can call those functions from it. Additionally, other developers can easily call these functions in their pipeline. Currently, nine packages in total are included to perform the whole workflow, from raw data processing to biological function mining, and the many graphic functions allow users to generate publication-quality graphics (https://massdataset.tidymass.org/articles/ggplot_mass_dataset). Most of the functions and tools that are widely used are included or supported in tidyMass. For functions/tools that are not yet wrapped, it is simple to implement and integrate them with tidyMass. Finally, one package named “tidymass” was developed to easily install and manage all the packages in the project. For each package, a website with function and package-level help documents and reproducible examples was created to guide new users on how to use it.

#### Naming and Coding style

In tidyMass, we strove to provide concise and meaningful names. To make the tidymass more user-friendly and easier to use, the coding style of tidyMass follows the tidyverse style guide (https://style.tidyverse.org/). Briefly, all the names of packages in the tidyMass project start from “mass” or “met” and follow a noun to describe their function. Such as “massCleaner” which is used for data cleaning and “massQC” which is used for data quality assessment. The variable and function names follow the snake case naming policy, using only lowercase letters, numbers, and underscores are used to separate words within a name. Generally, all the variable names are nouns, and function names are verbs.

#### Help document and tutorials

We provide the function-level, package-level, and pipeline-level help documents and tutorials as a learning guide for tidyMass. For the function-level help document, the users can find it on the “Reference” page on the corresponding website for each package. It is also possible to access them quickly in the R environment using the “?” function. For the package-level and pipeline-level, the websites are created using the “pkgdown” tools for all the packages, the users can find the help document or tutorial on the “Help document” or “Tutorial” page.

#### Do not reinvent the wheel

When we designed tidyMass, another important rule was that we did not want to create redundant tools which have similar functions with existing tools. For example, when we want to remove variables from “mass_dataset”, the “filter()” function from the dplyr package is more efficient and popular in the R community. We do not need to create a new function to process the “removing features” step. So in tidyMass, we made use of the base or popular functions in R to support “mass_dataset” to operate the same functions (https://massdataset.tidymass.org/articles/tidyverse_verse, https://massdataset.tidymass.org/articles/base_function). This design also makes it easier for new users to adopt tidyMass and reduce their study burden, it also means that the tidyMass code is more readable and shareable.

#### Deployment and Installation

All the packages in the tidyMass project are open-source and can be accessed publicly. In case the internet is not stable for one code hosting platform, we deployed it in three different code hosting platforms, namely GitHub (https://github.com/tidymass), GitLab (https://gitlab.com/dashboard/projects), and Gitee (https://gitee.com/jaspershen/dashboard/projects). Any changes will be updated on the three platforms at the same time, so the users can access and install them from at least one platform in any situation.

### MassDataset package

The massDataset package is used to provide a uniform data form/structure for LC-MS-based untargeted metabolomics data, relevant metadata, and the corresponding processing parameters (https://massdataset.tidymass.org/). Several packages in R provide the object-oriented class for efficient manipulation of sequencing data^14,20^, and although a similar concept (XCMS3, https://github.com/sneumann/xcms) has been also utilized in the metabolomics field^18^, there is still no specific uniform data form for all the processing/analysis workflow for LC-MS-based untargeted metabolomics data. Therefore, the massDataset package, the “mass_dataset” class, was specifically designed to store and manage processed metabolomics data and represents this data as an instance of the main data class. This is a key feature of the tidyMass project, all the subsequent wrapped operation functions use this class as their sole or primary input data form.

#### The “mass_dataset” class

The “mass_dataset” is an S4 object in the R environment that contains nine components (**Fig. 2**), including 1) expression data (expression_data) is a data frame that represents the abundance of all the metabolic features (peaks) in all samples. Each row is a metabolic feature (peak) and each column is a sample. 2) Sample information (sample_info) is a data frame that represents the metadata of samples. The first column is the sample IDs which should be completely identical to the column names of the expression data. Other columns are the attributes of samples, such as subject ID, sample batch, injection order, *etc.* 3) Variable information (variable_info) is a data frame that represents the metadata of variables (metabolic features or peaks). The first column is the variable IDs which should be completely identical to the row names of the expression data. Other columns are the attributes of variables, such as m/z, rt and mean intensity, *etc.* 4) Variable information note (variable_info_note) is a data frame that represents the metadata of variable information. 5) Sample information note (sample_info_note) is a data frame that represents the metadata of sample information. 6) MS^2^ spectra (ms2_data) is a list (“ms2_data” class) that is used to store the MS^2^ spectra for peaks. For each spectrum, the parent ion information, MS^2^ spectrum (a data frame with fragment ion m/z and intensity), and the corresponding peak are stored. 7) Annotation result (annotation_table) is a data frame representing the annotation results for variables. 8) Processing information (process_info) is a list (“tidymass_parameters” class) that is used to store the parameters for each processing/analysis step that has been applied on the “mass_dataset” class.

#### Automatic synchronization of components in the “mass_dataset” class

The components in “mass_dataset” are relevant. For example, the columns of expression data should completely correspond to the rows of sample information, and the rows of expression data should completely correspond to the rows of variable information. When one component in the “mass_dataset” class is modified, other components which are relevant to the changed component will automatically change to keep the consistency of all the components (**Fig. S2**). This design makes it easier to modify the datasets and keep them consistent.

#### The addition of MS^2^ data to the “mass_dataset” class

MS^2^ spectra data is important for LC-MS-based untargeted metabolomics data for metabolite annotation. One MS^2^ spectrum is defined by the spectrum information (parent ion information) and MS^2^ spectrum data frame. The spectrum information records the parent ion m/z, retention time, and other information. The MS^2^ spectrum data frame is a matrix with two columns, fragment *m/z,* and intensity. In the massDataset package, the MS^2^ spectra data can be added to the “mass_dataset” class using the “mutate_ms2()” function. Briefly, the MS^2^ spectra are extracted from the MS^2^ data files (mgf format), and then for each MS^2^ spectrum, it will be assigned to metabolic features based on m/z and retention time matching^21^. To organize and process the MS^2^ data in the “mass_dataset” class, a class named “ms2_data” is designed.

#### Base operation functions for the “mass_dataset” class

Base operation functions have been provided in the massDataset package to process the “mass_dataset” class. The functions can be divided into four classes (**Fig. S3**). 1) The first class of functions is used to extract and output datasets in “mass_dataset”. 2) The second class of functions is used to summarize and explore data. 3) The third class of functions is used to preprocess data. For example, add new information, remove samples/variables. 4) The fourth class functions are used to combine or merge two “mass_dataset” class objects. To reduce the difficulty and cost of learning, for the functions which are widely used in R for the same aims but other objects, we wrapped them in massDataset and made the “mass_dataset” class as their input data form. For example, the “filter()” functions from the tidyverse package are widely used in data science to remove eligible variables, so this function is wrapped and users can filter variables from any components in “mass_dataset”.

#### The “tidymass_parameter” class

To store the parameters for each step that is applied on the “mass_dataset” class, a *“*tidymass_parameter*”* class was designed in massDataset. Briefly, four slots are in the “tidymass_parameter” class, namely package name, function name, processing time, and parameter list. The parameter is stored as a list, whose items are specific settings, and the names are arguments. The “tidymass_parameter” classes for all the processing/analysis steps are stored in the “process_info” slot of the “mass_dataset” class and ordered by processing time. Thus, it is possible and easy for the users to trace the processing and analysis for this object. This is another key design in the tidyMass project, which provides the fundamentals for reproducible analysis.

### MassConverter package

The massConverter package is used to convert mass spectrometry raw data to different format data (https://massconverter.tidymass.org/). MSconvertGUI is the interactive version of the msconvert tool for converting mass spec data files to various formats, which is widely used in the metabolomics field^8^. It also provides the command line version. However, it is software that can only be installed on Windows OS, so cannot be used by Mac OS and Linux users. To achieve a comprehensive reproducible analysis, it is important to do the data converting and record the parameters in the R environment.

#### Docker version of msconvert

The team provides a docker version of msconvert (pwiz, https://hub.docker.com/r/chambm/pwiz-skyline-i-agree-to-the-vendor-licenses), so the massConverter package can convert mass spectrometry raw data to different formats. The users need to install docker based on the official website (https://www.docker.com/get-started). Then they pull the pwiz image by using the “docker_pull_pwiz()” function, which will download the pwiz image from the docker hub, therefore it can be used for converting data.

#### Convert data

Many parameters are included in the mass spectrometry data conversion. The “create_msconvert_parameter()” function is used to set the converting parameters. The detailed converting parameters and their meanings can be found in **Table S2**. After setting the parameters, the “convert_raw_data()” function is used to convert the raw data to other formats. The massConverter package makes it possible to convert the mass spectrometry data using R, and integrate data converting steps with other data processing and analysis in one code file, making the reproducible analysis of metabolomics data more efficient.

### MassProcesser package

The massProcesser package is used for mass spectrometry raw data processing, including peak picking and peak grouping based on the widely used XCMS^9^ (https://massprocesser.tidymass.org/). We have added some new functions to make the results more interpretable. After the processing, a “mass_dataset” class is generated with simple sample information. Then users can add more information directly to it for subsequent processing and analysis using other packages from the tidyMass project. This makes it smoother and more straightforward to combine raw data processing and other processing/analysis steps. In addition, all the graphics from massProcesser, such as “BPC”, “TIC”, and “retention time correction” are generated using the ggplot2 package, which generates high-quality figures for publication. Another important feature of the massProcesser package is that the users can easily extract and score the EIC of all the features and evaluate the quality of features, so can avoid false-positive findings in the subsequent analysis. The raw data processing is optional in the whole workflow, users can use other software/tools to generate peak tables.

### MassCleaner package

The massCleaner package is used to do the data cleaning of metabolomics data (https://masscleaner.tidymass.org/). The LC-MS-based untargeted metabolomics data always contain different types of bias arising from sample preparation and data acquisition (e.g., contamination, drift in signal intensity.), this is the reason to perform data cleaning as an essential step, which is used to remove unwanted variations. It can be divided into different steps, and some steps are optional, and the orders can be customized based on the study design and aims.

#### Noisy feature removal

The noisy feature removal can be used based on different rules, according to the experimental aims and design. The functions in massDataset and other packages make it simple to perform the noisy feature removal. For example, the users can define the noisy features as the metabolic features that have missing values more than in 20% QC samples or in 50% subject samples. So the “mutate_variable_na_freq()” function can be added to variable information and then remove the noisy features using the “filter()” function from the dplyr package.

#### Outlier samples removal

Outlier samples are a recurrent problem, especially when analyzing large cohorts. Detecting and removing the outlier samples are critical to avoid false positive and false negative findings in the subsequent analysis. Different methods have been used to define and detect outlier samples in tidyMass^22^. The first rule is the missing value percentage for each sample^10^. If one sample with more than 50% features is missing values, it means that there may be issues in the sample preparation or data acquisition, so those samples are labeled as an outlier. Other methods are also included to detect outlier samples^23^. In brief, all the biological subject samples are used for PCA analysis, then the samples whose principal component 1 (PC1) are more than 6 standard deviations away from the mean value will be labeled as outlier samples. To make this method more robust, we also calculate the median instead of the mean value, and MAD (median absolute deviation) instead of the SD (standard variation) because they are more robust estimators. The last method is based on distance. Instead of using the infinite distance, Mahalanobis distance is a multivariate distance based on all variables (principal components) at once. We use a robust version of this distance, which is implemented in packages robust and “robustbase” and that is reexported in “bigutilsr”. Once the outliers have been detected by different methods, it is easy for users to remove the samples from the “mass_dataset” class according to their study aims using the “filter()” function.

#### Missing value imputation

Missing value imputation should be performed after noisy features and outlier sample removal. In massCleaner, four widely used methods are implemented to perform missing value imputation: 1) K-nearest neighbors (KNN)^24^, 2) Bayesian principal component analysis replacement (BPCA)^25^, 3) svdImpute^26^, 4) random forest imputation (missForest)^27^, 5) zero values, 6) mean values, 7) median values and 8) minimum values. KNN is recommended to impute missing values and set them as default^10^.

#### Data normalization and integration

Data normalization and integration are important to remove the unwanted analytical variations occurring in intra- and inter-batch measurements and to integrate multiple batches forming an integral data set for subsequent statistical analysis^28^. In the metCleaner package, several methods that are widely used are integrated. The methods can be divided into two different classes. The first class is the sample-wise method, including PQN, median, mean, total intensity normalization^12^. Total intensity normalization means that all the variable intensity is divided by the total intensity of all the variables in one sample. This method sets the total sum of signals to a constant value for each sample. The median and mean normalization have the same concept. However, these approaches could be hampered. For instance, in the case of large mass differences between samples that may lead to different variable extraction efficiencies between samples^12^. The second class is the QC sample-based data normalization, including SVR^29^, and LOESS^3^. QC samples are typically generated by mixing aliquots of each subject sample and are regularly analyzed during an experimental run to monitor the stability of the analytical platform and are particularly useful for identifying batch effects. For QC-based data normalization, they require that the first and last injections should be QC samples. The data integration method is used to integrate multiple batch data. In the massCleaner function, the QC median, QC mean, subject means, the subject median for each variable (metabolic feature or peak) can be used as the correction factors to integrate batches^29^.

### MassQC package

The massQC package is used to assess the data quality of LC-MS-based untargeted metabolomics (https://massqc.tidymass.org/). The data quality of metabolomics is visually assessed by several aspects^10^. 1) Missing value distribution across samples and/or variables. If one variable (metabolic feature or peak) has more missing values, it means that this variable may be a noisy feature. The same applies to samples that have lots of missing values, it could signify that they are outlier samples that should be removed. 2) RSD (relative standard deviation) for all variables in QC (quality control) samples. Since the QC samples are similar and injected frequently during the data acquisition, the RSD of variables in QC samples can be utilized to evaluate the stability of LC and mass spectrometry. In biomarker discovery, the cutoff is always set as 30%. 3) Intensity of all variables in samples. For QC samples, the median value of the intensity of all variables should be very close. 4) The correlation of QC samples. 5) PCA score plot. The high-quality data assessed by a PCA should show a tight clustering of QC samples relative to the distribution of non-QC samples. In massQC, the users can use the “mass_dataset” as the argument to get the result for each aspect in any step of the whole workflow. In addition, one function named “massqc_report()” can be used to generate an HTML format report including all the results, which is very convenient (https://massqc.tidymass.org/articles/html_qa_report).

### MetID package

The metID package is used to perform metabolite annotation based on in-house and available open-source databases (https://metid.tidymass.org/)^11^. It combines information from all major databases for comprehensive and streamlined compound annotation. MetID is a flexible, simple, and powerful tool allowing the compound annotation process to be fully automatic and reproducible. What should be noted is that metID^11^ was not originally designed for the tidyMass project, so it doesn’t support “mass_dataset”. However, it is simple to integrate with tidyMass, which demonstrates the flexibility and extensibility of tidyMass. To integrate metID with the tidyMass project, a function named “annotate_metabolites_mass_dataset()” has been developed to support the “mass_dataset” class. All the annotation results have been organized as a data frame and assigned to “annotation_table” in the “mass_dataset” class. The annotation parameters (matching parameters, the database used, *etc.*) are also assigned to processing information. The users can access the annotation table in “mass_dataset” by using the “extract_annotation_result()” function.

### MassStat package

The massStat package is used to perform common statistical analyses within metabolomics analysis (https://massstat.tidymass.org/). The massStat package provides efficient tools for the different steps required within the complete data analytics workflow: scaling, univariate analysis, multiple testing correction, multivariate analysis, candidate biomarkers selection, and correlation network analysis.

#### Scaling

Scaling is a procedure where each variable is modified by a factor and accounts for the different statistical characteristics of each variable. Without scaling, highly abundant compounds tend to dominate the analysis when variance-dependent techniques such as PCA are used. Now in massStat, three commonly used scaling methods are included. Unit-variance scaling (uv) divides each variable by its standard deviation. Pareto scaling, intermediate between no scaling and uv scaling, divides each variable by the square root of the standard deviation. Range scaling divides each variable by its range in all the samples.

#### Univariate analysis

Commonly used univariate analysis tools have been implemented in tidyMass. Student's t-test (t.test), and Wilcoxon signed-rank test (wilcox.test). The different multiple testing correction methods from p.adjust are also implemented. The fold change, p values, and adjusted p values are directly added to the variable information in “mass_dataset”.

#### Correlation, distance, and correlation network

Correlation and distance between samples or variables can be calculated using the “cor_mass_dataset()” and “dist_mass_dataset()” functions. The “margin” argument is provided in both functions which requests the sample or variable correlation/distance matrix. The correlation network is widely used to explore the co-expression and co-regulation metabolites, in massStat, the users can obtain a network data format (from ggraph and tidygraph packages) from the “mass_dataset” class object. Then this object can be used for network analysis and visualization using the powerful network analysis ecosystem, including ggraph, igrpah, and tidygraph.

#### Multivariate analysis

It is possible to perform various multivariate analyses, such as PCA, PLS, PLS-DA. A typical first-pass unsupervised method used in untargeted LC-MS-based metabolomics is PCA. The score scatter plots, where each sample is depicted as a point, reveal how all samples relate to each other. Supervised methods such as PLS, PLS-DA^30^, and clustering are also provided.

### MetPath package

The metPath package enables pathway enrichment analysis for metabolomics (https://metpath.tidymass.org/). At present, metPath provides two commonly used metabolic pathways for this analysis, KEGG^31^, and SMPDB^32^. To organize and manage the pathway database, a class named “pathway_database” was designed in the metPath package, which is used to store and manage the pathway data. Like the “mass_dataset” class, the “pathway_database” class can be operated by the base and tidyverse functions, which makes it easy to process and manage the pathway database. Then the Hypergeometric test or Fisher's exact test is performed for pathway enrichment. Different visualization methods for enriched pathways are also provided based on ggplot2 to generate high-quality graphics.

### MassTools package

The massTools package provides useful tiny functions for mass spectrometry data processing and analysis (https://masstools.tidymass.org/). It is a supporting and base package for the tidyMass project. Some functions are universal and may be used and called by different packages, so they are placed in the massTools package, therefore other packages can directly call those functions anytime and anywhere. For example, the MS^2^ spectra matching plot can be used in different places, so it is also placed in the massTools package.

### TidyMass package

The tidyMass package is designed to organize and manage all the packages in the tidyMass project (https://tidymass.tidymass.org/), allowing for easy installation and loading multiple “tidyMass” packages in a single step. In brief, all the other packages in the tidyMass project are set as the dependent packages of it, so the users can install all the packages in the tidyMass project by only installing the tidymass package. When one or more packages are updated in the tidyMass project, then users can easily check and update them using the tidyMass package. In addition, users can load all the packages into the R environment by only loading the tidyMass package.

### Extend tidyMass project

An increasing number of data processing and analysis tools are being developed within the field of metabolomics. This could be problematic, as the integration of these functions and tools is needed to enable their use in tidyMass. However, the specific and uniform data form (“mass_dataset” class) simplifies the integration of tools that are not wrapped in tidyMass for developers. In fact, in tidyMass, the R base function, tidyverse, and metID package have been integrated with the “mass_dataset” class. In brief, the function should change the “mass_dataset” as its supporting object, and then call the function to process or analyze. A protocol is available to show how to make a function that supports the “mass_dataset” class (https://massdataset.tidymass.org/articles/based_on_mass_dataset). In addition, it is easy to integrate tidyMass with other pipelines. For example, xcmsrocker is an open-source project (https://github.com/yufree/xcmsrocker) which was created and maintained by Dr. Miao Yu, this project houses various R packages for LC-MS-based metabolomics data processing and analysis, and tidyMass was recently implemented into this project. Another example is the Stanford Data Ocean (https://innovations.stanford.edu/sdo), which is a cloud-based computation platform for multi-omics data processing and analysis, and tidyMass is also implemented onto it.

### Data preparation for tidyMass

TidyMass is a flexible pipeline that utilizes the modular design concept, which means that the user can perform a comprehensive and full data processing workflow for metabolomics or can choose to perform various or multiple steps of the workflow.

#### Data preparation for massProcesser

If the users use the massProcesser package for raw data processing, the mzXML (or mzML) data format should be prepared. All the mzXML format files should be placed in different folders according to their class or group. For example, QC samples and blank samples should be placed into folders named “QC” and “Blank” folders, respectively. Biological subject samples can be placed in a folder named “Subject” or placed into different folders that are named according to the class of samples, for example, “Control” or “Case”.

#### Data preparation for other packages

The users can also use other software to perform raw data processing to generate the peak (metabolic feature) table, such as MS-DIAL^32,33^, mzMine^34^, *etc*. Then the data can be prepared and the “create_mass_dataset()” function is used to generate the “mass_dataset” class object. These files are required for the “create_mass_dataset()” class. The first file is “expression_data” which is a matrix to store the abundance for each variable in each sample. The column is a sample, and the row is variable. The second file is “sample_info” which is a matrix to store the metadata of samples. What should be noted is that the first column is sample ID (sample_id) which is completely identical to the column names of expression data. The third file is “variable_info” which is a matrix to store the metadata of variables. The first column is the variable ID (variable_id) which should be completely identical to the row names of expression data. In addition, the second column and third column should be mass-to-charge ratio (m/z) and retention time (rt, the unit is second), respectively, which are specific spectral information for mass spectrometry data.

### Reproducible analysis using tidyMass

One of the most important aims of tidyMass is to improve the reproducible analysis of LC-MS-based untargeted metabolomics data. In tidyMass, the “mass_dataset” class and modular design make it easier for data sharing and reproducible analysis for metabolomics data.

#### Data sharing

We have enabled a straightforward method for tidyMass users to share their processed data. After preparing the datasets, a “mass_dataset” class object can be generated using the massDataset package, and then users can share the “mass_dataset” class object with collaborators without the need to share multiple files, which is the typical way of sharing this type of data. Collaborators can load the shared “mass_dataset” class object in the R environment and then directly and easily process it using tidyMass. The users can also output all the components in the “mass_dataset” class to xlsx or csv format, and share one or several files of their choosing.

#### Reproducible analysis

We encourage users to share their data (“mass_dataset” class) and tidyMass pipeline with other collaborators or journals using R script or R markdown files. As the data processing and analysis code is written by R (tidyMass pipeline), it is straightforward for collaborators to easily reproduce the analysis and results. The demo data (“mass_dataset” class) and R code (R markdown) for our demo data have been provided on the tidyMass homepage (https://www.tidymass.org/start/). The demo data and R script of the case study presented are also downloadable on the homepage (https://www.tidymass.org/start/demo_data/).

#### Docker image of tidyMass

A docker image of tidyMass named “tidymass” has been deployed on the docker hub (https://hub.docker.com/r/jaspershen/tidymass). This docker image was developed based on the rocker image verse (https://hub.docker.com/r/rocker/verse), which contains a Rstudio and R environment, and installed most of the widely used data science packages, such as tidyverse. We installed all the packages in tidyMass with associated dependent packages, the demo datasets and code were also implemented. The new docker image was then built named “tidymass”. The docker version of tidyMass can be used for data analysis by downloading it and then opening the website version Rstuido for data analysis. The “tidymass” image can also be used as a base image for users who want to build a new image to share their analysis environment with other collaborators or reviewers to repeat their analysis and results. A protocol on how to use the docker image of tidyMass is provided on the website of tidyMass (https://www.tidymass.org/start/tidymass_docker/).

### Sample preparation and analytical conditions for the case study

All the sample preparation and analytical conditions for the case study can be found in our previous publication^15^.

## Data availability

All the demo data for how to use tidyMass can be accessible on the tidyMass website (https://www.tidymass.org/). For the case study, mass spectrometry raw converted data (mzML) for the case study in this paper is accessible on MetaboLights with MTBLS1122 (HILIC positive), MTBLS1124 (HILIC negative), MTBLS1122 (RPLC positive) and MTBLS1130 (RPLC negative). The MS^2^ data (mgf) and processed data (“mass_dataset” class) from the massProcesser package are available on the tidyMass project website (https://www.tidymass.org/start/case_study/), and the “mass_dataset” objects are provided as **Supplementary Data 1**.

## Code availability

All the source code of the tidyMass project is deployed on GitHub (https://github.com/tidymass), GitLab (https://gitlab.com/users/jaspershen/projects), and Gitee (https://gitee.com/jaspershen/projects), and are public under the MIT License; and works on Windows, macOS X, and most Linux distributions. The docker image of tidyMass is hosted on the docker hub (https://hub.docker.com/r/jaspershen/tidymass). The code of the case study (Rmarkdown format, https://www.tidymass.org/start/case_study/) is provided as **Supplementary Data 2**.

## Acknowledgments

We thank Dr. Miao Yu for integrating the tidyMass project with the xcmsrocker project and providing advice on the development of the docker image version of tidyMass. We also thank Dr. Amir Bahmani and Kexin Cha for integrating the tidyMass project with the Stanford Data Ocean project. We also thank Dr. Axel Brunger for the advice on the manuscript. C.H.J was supported by the NCI/NIH under Award Number K12CA215110, NIGMS/NIH under Award Number 1RM1GM141649-01, and American Cancer Society Research Scholar Grant 134273-RSG-20-065-01-TBE.

## Author contributions

X.S. and M.P.S. conceived the method and supervised its implementation. X.S. developed the methods, packages, and the docker image. X.S. and C.W. built the websites and wrote the help documents and tutorials. H.Y. provided and prepared the case study data, H.Y. and X.S. analyzed the case study data. X.S., H.Y., and C.W. prepared the figures. X.S, H.Y., C.W., C.H.J, and M.S.P wrote the manuscript, C.H.J, M.S.P, and P.G. improved the manuscript. All authors contributed to the final manuscript.

## Additional information

**Correspondence and requests for materials** should be addressed to X.S. or M.P.S.

