## Supplementary for "*TidyMass*: An Object-oriented Reproducible Analysis Framework for LC-MS Data"

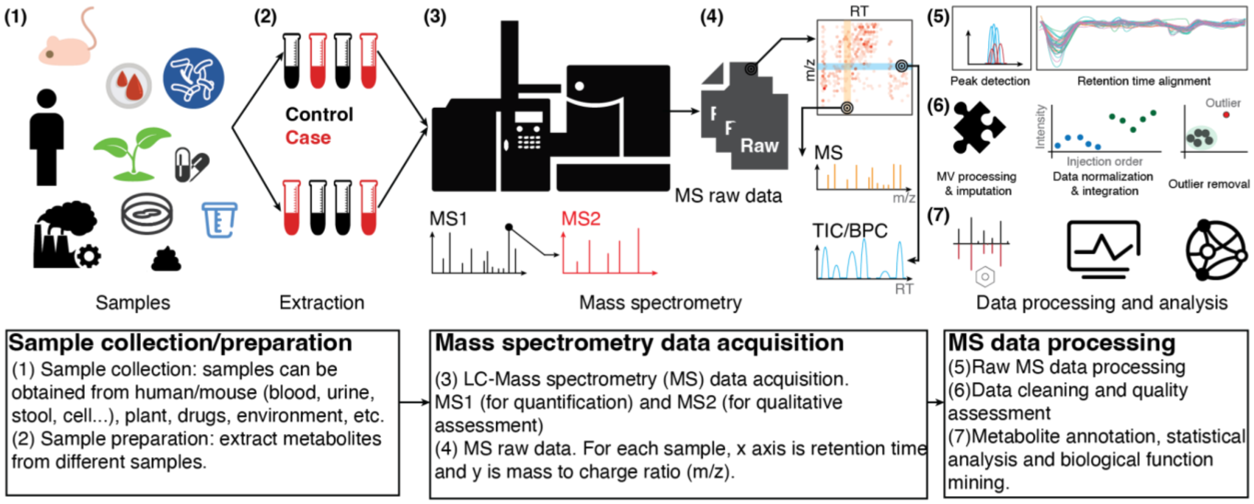


**Supplementary Figure 1. Example of an untargeted high-resolution LC-MS experiment.** A diagram of an experimental and analysis workflow for LC-MS-based untargeted metabolomics. LC-MS-based untargeted metabolomics involves several fundamental steps: **(1)** sample collection and preparation; **(2)** metabolite extraction; **(3)** mass spectrometry data acquisition and raw data generation; **(4)**, **(5)** raw data processing; **(6)** data cleaning; and **(7)** metabolite annotation and biological function mining. The intended role for *tidymass* is in MS data processing and analysis (steps 5-7).

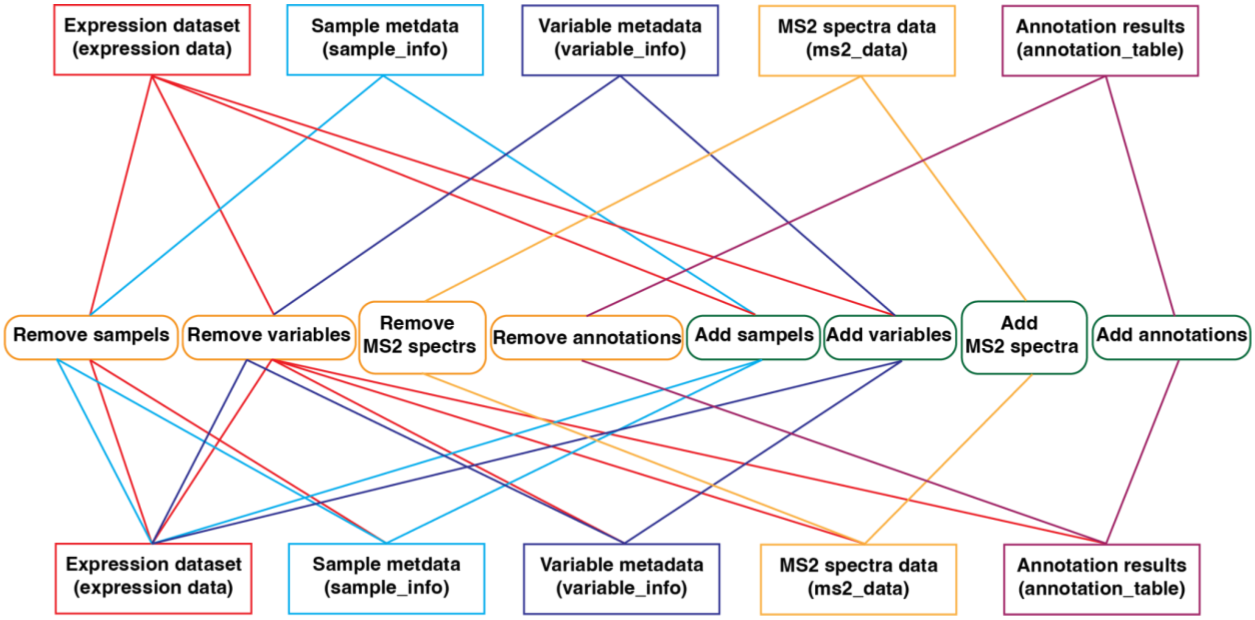


**Supplementary Figure 2. Automatic synchronization of components in the "mass_dataset" class.**

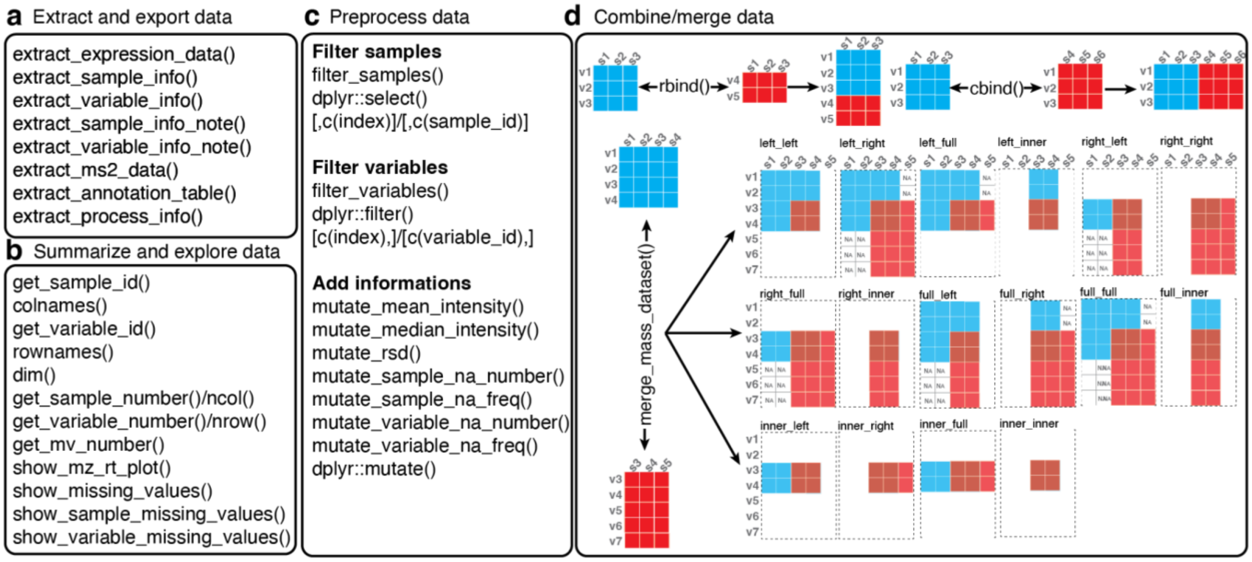


**Supplementary Figure 3. Functions that support the "mass_dataset" class. (a)** Functions to extract and export data. **(b)** Functions to summarize and explore data. **(c)** Functions for preprocessing data. (d) Functions to combine/merge data.

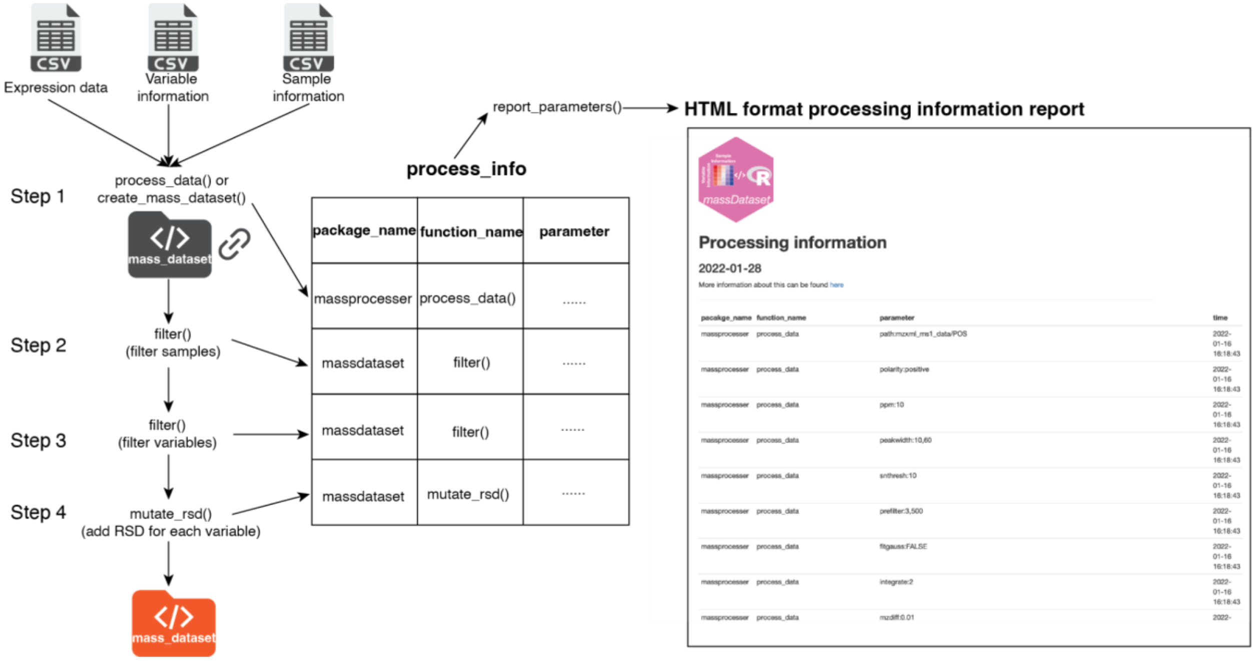


**Supplementary Figure 4. Processing information in the "mass_dataset" class.** Different functions from different packages can apply to the "mass_dataset" class step by step, and the parameters will be recorded in the "process_info" slot. The "report_parameters()" function from the massDataset package can be used to extract them and output them as an HTML format processing information report.

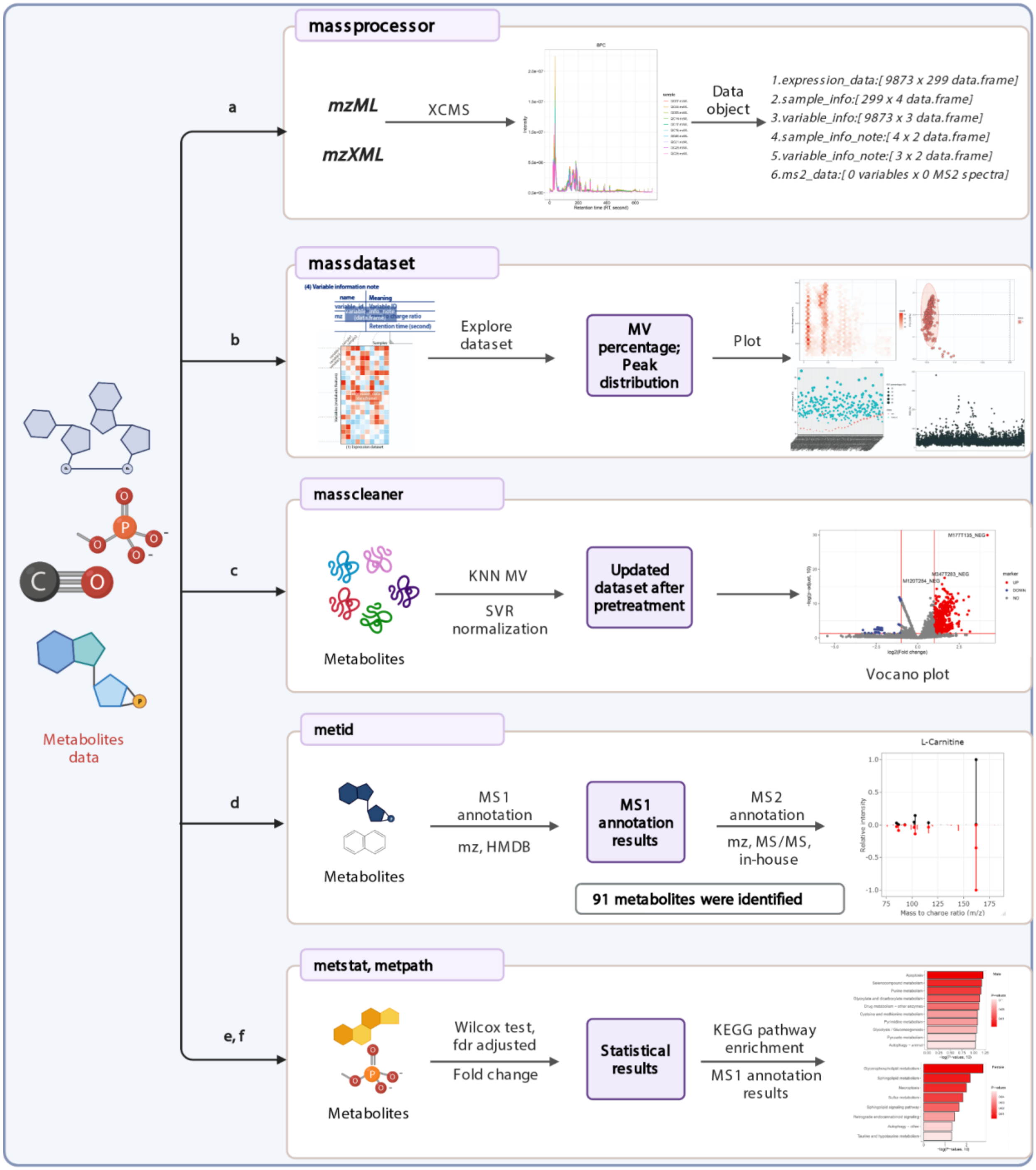


**Supplementary Figure 5. Analysis workflow of the case study.** **(a)**, **(b)**, **(c)**, **(d)**, **(e)**, and **(f)** correspond to the step in **Fig. 2**.

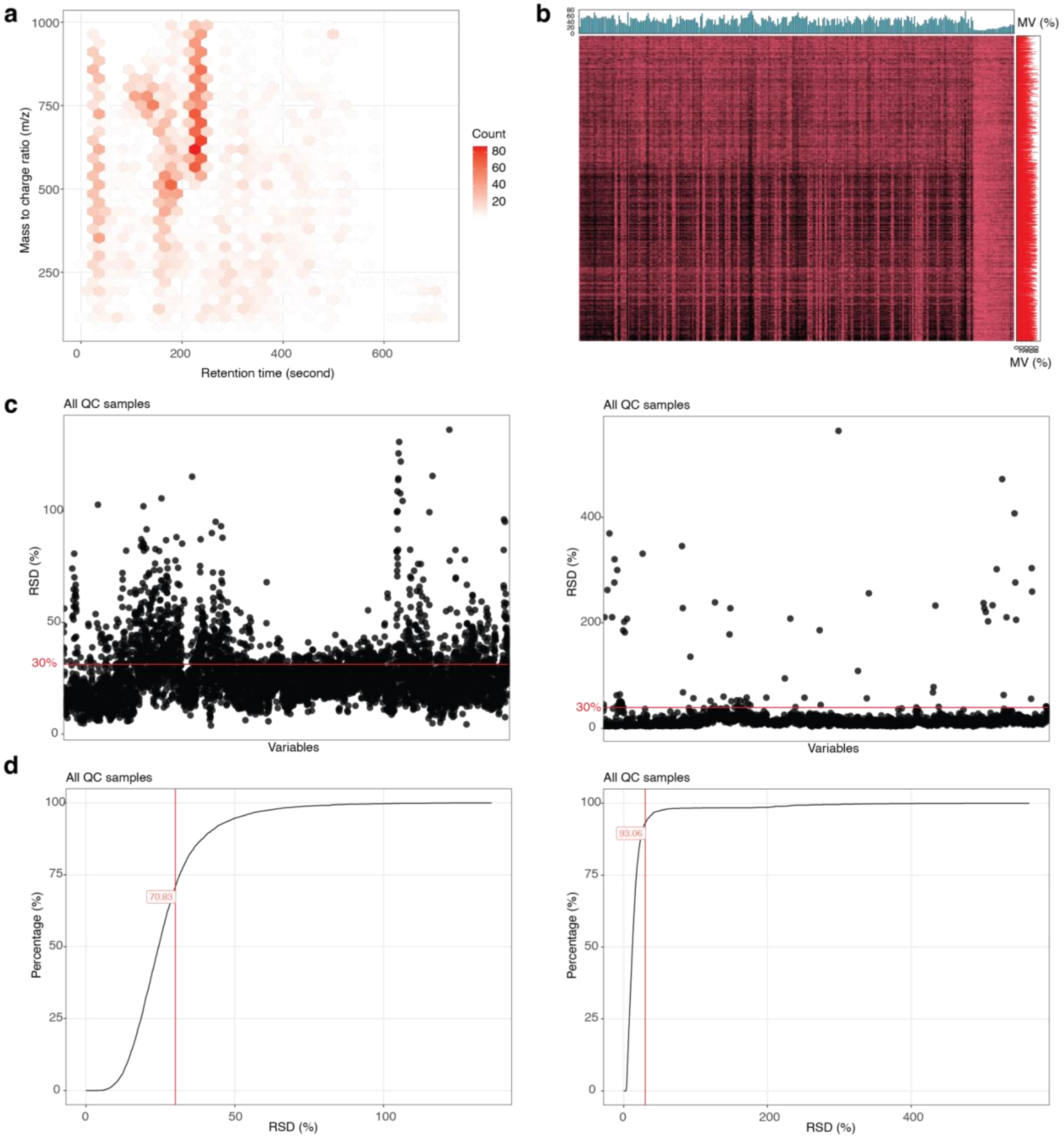


**Supplementary Figure 6. Data quality assessment before and after data cleaning using tidyMass.** Here the HILIC positive mode data is used as an example. **(a)** The distribution of metabolic features. **(b)** Missing distribution. The X-axis displays samples and the y-axis displays metabolic features. **(c)** RSD for all the metabolic features in QC samples before (left) and after data cleaning (right). **(d)** RSD cumulative plot before (left) and after data cleaning (right).

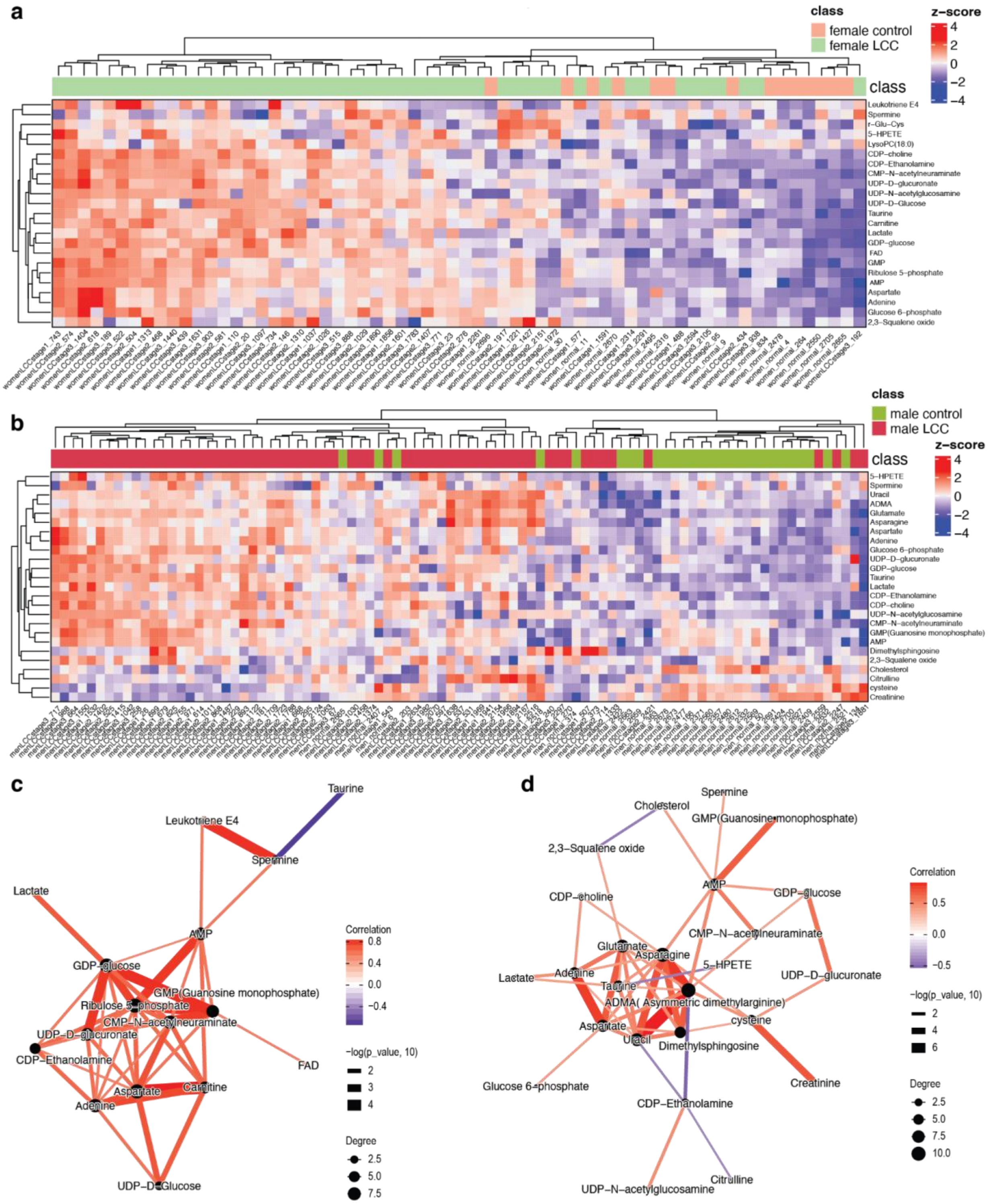


**Supplementary Figure 7. Differential expressed metabolites (DEMs) for tumors from female and male patients with left-sided colorectal cancer (LCC). (a)** Heatmap for normal colon (control) and tumor tissues from female patients using DEMs. **(b)** Heatmap for normal colon (control) and tumor tissues from male patients using DEMs. **(c)** Correlation networks for metabolites dysregulated in tumors compared to normal tissues from female **(c)** and male patients **(d)**.

**Supplementary Table 1.** Functions in the tidyMass project for data processing and analysis.

| Function name | package | Input data format | Output | Fcuntion |
|---|---|---|---|---|
| docker_pull_pwiz() | massConverter | - | - | Pull pwiz docker image. |
| create_msconvert_parameter() | massConverter | - | msconvert_parameter | Create parameter class for masssConverter. |
| convert_raw_data() | massConverter | Mass spectrometry raw data | Other format data | Convert MS raw data to other format data |
| create_mass_dataset() | massDataset | - | "mass_dataset" class object | Create "mass_dataset" class object |
| extract_expression_data() | massDataset | "mass_dataset" class object | data.frame | Extract expression data |
| extract_sample_info() | massDataset | "mass_dataset" class object | data.frame | Extract sample information |
| extract_variable_info() | massDataset | "mass_dataset" class object | data.frame | Extract variable inforamtion |
| extract_annotation_table() | massDataset | "mass_dataset" class object | data.frame | Extract annotation table |
| extract_variable_info_note() | massDataset | "mass_dataset" class object | data.frame | Extract variable information metadata |
| extract_sample_info_note() | massDataset | "mass_dataset" class object | data.frame | Extract sample information metadata |
| extract_process_info() | massDataset | "mass_dataset" class object | "tidymass_parameter" class | Extract processing information |
| extract_ms2_data() | massDataset | "mass_dataset" class object | "ms2_data" class | Extarc $MS^2$ data |
| mz_rt_match() | massTools | "mass_dataset" class object | data.frame | Match two "mass_dataset" class according to mz and rt |
| fliter_samples() | massDataset | "mass_dataset" class object | "mass_dataset" class object | Filter samples |
| filter_variabels() | massDataset | "mass_dataset" class object | "mass_dataset" class object | Filter variables |

| | | | | |
|---|---|---|---|---|
| filter() | dplyr | “mass_dataset” class object | “mass_dataset” class object | Filter rows from components in “mass_dataset” class |
| mutate() | dplyr | “mass_dataset” class object | “mass_dataset” class object | Mutate new columns to components in “mass_dataset” class |
| select() | dplyr | “mass_dataset” class object | “mass_dataset” class object | Select columns from components in “mass_dataset” class |
| mutate_ms2() | massDataset | “mass_dataset” class object | “mass_dataset” class object | Add $MS^2$ data to “mass_dataset” class |
| mutate_mean_intensity() | massDataset | “mass_dataset” class object | “mass_dataset” class object | Add mean intensity to variable information |
| mutate_median_intensity() | massDataset | “mass_dataset” class object | “mass_dataset” class object | Add median intensity to variabel information |
| mutate_rsd() | massDataset | “mass_dataset” class object | “mass_dataset” class object | Add RSD to variable information |
| mutate_sample_na_number() | massDataset | “mass_dataset” class object | “mass_dataset” class object | Add variable NA numbers to sample inforamtion |
| mutate_sample_na_freq() | massDataset | “mass_dataset” class object | “mass_dataset” class object | Add variable NA frequency to sample inforamtion |
| mutate_sample_na_number() | massDataset | “mass_dataset” class object | “mass_dataset” class object | Add sample NA numbers to variable inforamtion |
| mutate_sample_na_freq() | massDataset | “mass_dataset” class object | “mass_dataset” class object | Add sample NA frequency to variable inforamtion |
| report_parameters() | massDataset | “mass_dataset” class object | HTML format report | Report for processing information |
| show_mz_rt_plot() | massDataset | “mass_dataset” class object | ggplot2 plot class object | Metabolic feature plot |
| show_missing_values() | massDataset | “mass_dataset” class object | ggplot2 plot class object | Missing value distribution plot |
| left_join() | dplyr | “mass_dataset” class object | “mass_dataset” class object | Mutate new columns to components in “mass_dataset” class |

| | | | | |
|---|---|---|---|---|
| cbind() | base | “mass_dataset” class object | “mass_dataset” class object | Bind two “mass_dataset” class objects by rows |
| rbind() | base | “mass_dataset” class object | “mass_dataset” class object | Bind two “mass_dataset” class objects by columns |
| merge_mass_dataset() | massDataset | “mass_dataset” class object | “mass_dataset” class object | Merge two “mass_dataset” class objects |
| process_data() | massProcesser | mzXML format data | “mass_dataset” class object | Raw data processing |
| extract_eic() | massProcesser | - | - | Extract EIC |
| detect_outlier() | massCleaner | “mass_dataset” class object | “outlier_samples” class object | Detect outlier samples |
| extract_outlier_table() | massCleaner | “outlier_samples” class object | data.frame | Outlier sample table |
| impute_mv() | massCleaner | “mass_dataset” class object | “mass_dataset” class object | Impute missing values |
| integrate_data() | massCleaner | “mass_dataset” class object | “mass_dataset” class object | Interate data |
| normalize_data() | massCleaner | “mass_dataset” class object | “mass_dataset” class object | Normalize data |
| optimize_loess_span() | massCleaner | “mass_dataset” class object | data.frame | Optimize the parameters for loess regression |
| align_batch() | massCleaner | “mass_dataset” class object | “mass_dataset” class object | Align two “mass_dataset” class objects |
| massqc_cumulative_rsd_plot() | massQC | “mass_dataset” class object | ggplot2 plot class object | Cummulative RSD plot |
| massqc_pca() | massQC | “mass_dataset” class object | ggplot2 plot class object | PCA score plot |
| massqc_rsd_plot() | massQC | “mass_dataset” class object | ggplot2 plot class object | RSD distribution plot |
| massqc_sample_boxplot() | massQC | “mass_dataset” class object | ggplot2 plot class object | Sample box plot |
| massqc_sample_correlation() | massQC | “mass_dataset” class object | ggplot2 plot class object | Smaple correlation plot |
| massqc_report() | massQC | “mass_dataset” class | HTML format report | HTML format QC |

| | | | | |
|---|---|---|---|---|
| | | object | | report |
| annotate_metabolites_mass_dataset() | metID | “mass_dataset” class object | “mass_dataset” class object | Annotate features in “mass_dataset” by databases |
| annotate_single_peak_mass_dataset() | metID | “mass_dataset” class object | “mass_dataset” class object | Annotate one feature in “mass_dataset” by databases |
| ms2_plot_mass_dataset() | metID | “mass_dataset” class object | ggplot2 plot class object | $MS^2$ matching plot |
| convert_dummy_variable() | massStat | vector | data.frame | Convert vector to dummy variable |
| convert_mass_dataset2graph() | massStat | “mass_dataset” class object | tbl_gh class | Convert “mass_dataset” class to graph object |
| cor_mass_dataset() | massStat | “mass_dataset” class object | data.frame | Correlation matrix |
| dist_mass_dataset() | massStat | “mass_dataset” class object | data.frame | Distance matrix |
| Heatmap() | massStat | ComplexHeatmap | ggplot2 plot class object | Heatmap plot |
| pls() | massStat | mixOmics | “pls” object | PLS analysis |
| plsda() | massStat | mixOmics | “plsda” object | PLS-DA analysis |
| mutate_fc() | massStat | “mass_dataset” class object | “mass_dataset” class object | Add fold changs to variable inforamtion |
| mutate_p_value() | massStat | “mass_dataset” class object | “mass_dataset” class object | Univariable test |
| run_pca() | massStat | “mass_dataset” class object | pca class object | PCA analysis |
| pca_score_plot() | massStat | pca class object | ggplot2 plot class object | PCA score plot |
| scale_data() | massStat | “mass_dataset” class object | “mass_dataset” class object | Scale data |
| volcano_plot() | massStat | “mass_dataset” class | ggplot2 plot class object | Volcano plot |
| filter_pathway() | metPath | “pathway_database” class | “pathway_database” class | Filter pathways |
| get_hmdb_pathway() | metPath | - | “pathway_database” class | Get or download HMDB pathway |

| | | | | |
|---|---|---|---|---|
| get_kegg_pathway() | metPath | - | “pathway_database” class | Get or download KEGG pathway |
| enrich_kegg() | metPath | Query metabolite ID | “enrich_result” class | Pathway enrichment |
| enrich_hmdb() | metPath | Query metabolite ID | “enrich_result” class | Pathway enrichment |
| enrich_bar_plot() | metPath | “enrich_result” class | ggplot2 plot class object | Barplot to show the enriched pathways |
| enrich_scatter_plot() | metPath | “enrich_result” class | ggplot2 plot class object | Scatter plot to show the enriched pathways |
| enrich_network() | metPath | “enrich_result” class | ggplot2 plot class object | Network to show the enriched pathways |

**Supplementary Table 2.** Parameters for conversion using massConvereter.

| Argument name | Meaning | Format | Range |
|---|---|---|---|
| output_format | Output format | character | mzXML, mzXL, mz5, mgf, text, ms1, cms1, ms2, cms2 |
| binary_encoding_precision | Binary encoding precision | character | 32 or 64 |
| zlib = TRUE | Zlib or not | logical | TURE or FALSE |
| write_index | Write index or not | logical | TRUE or FALSE |
| peak_picking_algorithm | Peak picking algorithm | character | vendor, cwt, no |
| vendor_mslevels | Vender MS levels | vector | 1-n |
| cwt_mslevels | Cwt MS level | vector | 1-n |
| cwt_min_snr | Cwt minimum signal to noise rate | numeric | 0-1 |
| cwt_min_peak_spacing | Cwt minimum peak spacing | numeric | 0-1 |
| subset_polarity | Subset polarity | character | any, positive, negative |
| subset_scan_number | Subset scan number | vector | 0-n |
| subset_scan_time | Subset scan time | vector | 0-n |
| subset_mslevels | Subset MS level | vector | 1–n |
| zero_samples_mode | Zero sample mode | character | no, removeExtra, addMissing |
| zero_samples_mslevels | Zero sample MS levels | vector | 1-n |
| zero_samples_add_missing_flanking_zero_count | Zero sample add missing flanking zero count | numeric | 0-10 |

**Supplementary Note**

Colon tumor tissues and normal colon tissues were acquired from surgery and prospectively collected on 736 stage I-IV CRC patients in the period 1991–2001 at Memorial Sloan-Kettering Cancer Center (MSKCC, New York, NY, United States)[1]. Clinical data were recorded and updated retrospectively. Each tissue sample was snap-frozen in liquid nitrogen in the operating room and immediately stored in a −80 °C freezer. For this study, samples were selected that were from patients ≥ 55 years old to reduce the confounding effects of estrogen signaling on metabolism before menopause, also patients with early-onset CRC have a different etiology and biology than later-onset CRC. All normal colon tissues were selected from stage I-IV CRC patients (n = 39), and tumor tissue samples were selected from RCCs (right-sided colon cancer) and LCCs (left-sided colon cancer) stage I-III (n = 197). Only the normal and LCC samples were used for subsequent analysis. Clinical information was provided in **Table S3**.

**Supplementary Table 3.** Demographics of colon cancer patients from samples used within this study. Right-sided colon cancer = RCC, Left-sided colon cancer = LCC.

| | Normal (n=39) | Stage I (n=47) | | Stage II (n=86) | | Stage III (n=64) | |
|---|---|---|---|---|---|---|---|
| | | RCC (n=22) | LCC (n=25) | RCC (n=44) | LCC (n=42) | RCC (n=32) | LCC (n=32) |
| Sex, n | | | | | | | |
| Male | 27 | 10 | 15 | 23 | 25 | 15 | 14 |
| Female | 12 | 12 | 10 | 21 | 17 | 17 | 18 |
| Age, mean (sd) | | | | | | | |
| Male | 69.3 (9.6) | 73.9 (6.5) | 69.3 (5.8) | 72.9 (7.8) | 72.2 (8.5) | 73.5 (7.8) | 63.7 (5.8) |
| Female | 63.3 (16.1) | 72.1 (6.2) | 69.6 (7.6) | 73.5 (9.8) | 69.1 (7.8) | 72.2 (6.6) | 71.1 (6.0) |
| Race/Ethnicity, n | | | | | | | |
| NHWs | 35 | 20 | 22 | 40 | 37 | 26 | 25 |
| Hispanic | 2 | 2 | 3 | 2 | 3 | 4 | 2 |
| AA | 1 | 0 | 0 | 1 | 0 | 1 | 4 |
| API | 1 | 0 | 0 | 1 | 2 | 1 | 1 |

NHWs: non-Hispanic whites, AA: African-Americans, API: Asian-Pacific Islander

**Supplementary Data 1.** Processed data ("mass_dataset" class) from the massProcesser package.
**Supplementary Data 2.** Code file (Rmd format) for the case study.
